# Correlation/causation in ecology: towards mechanistic explanations for the Species-Area Relationship

**DOI:** 10.1101/044081

**Authors:** Andersonn Prestes

**Affiliations:** University of Hawaii at Manoa, Department of Plant and Environmental Protection Sciences. 3050 Maile Way, Gilmore 310. Honolulu, HI 96822, USA.

**Keywords:** Causation, Correlation, Species-richness, Diversification, Area, Pattern

## Abstract

There is a common intuition in biology that strict laws are very difficult to be found. Still, there are recurrent patterns in nature, suggesting broad generalizations and understanding of phenomena. The problem is that many generalizations in biology, especially in the form of correlations, might be decoupled from causality, weakening their power of explanation. Here, I bring an example on the Species-Area Relationship (SAR). The SAR is a well-known generalization in biology. The recurrent pattern states a positive relationship between area size and species richness. Understanding the mechanisms why there is a correlation between area and diversity remains a major challenge. I suggest an explicitly focus on mechanistic explanations for the SAR. I propose to use the integration, comparison and interpretation of other (associated or secondary) natural patterns in the searching for causal explanations. Area *per se* might not account for causality in species diversification or absolute species richness in larger regions. Biotic and abiotic factors of a given area might be studied in order to discover the causal underpinnings of the SAR.

## 1 Introduction

There is a common intuition in biology that strict laws are very difficult to be found. Usually, there are two arguments that favor the lack of strict laws: the complexity of biological systems (Rosenberg, 1994), and the nature of contingency in biological phenomena (Beatty, 1995). On the other hand, Sober (1997) and Elgin (2006) argue that al least some kind of lawlike generalizations in biology may hold true. They say that given a set of initial contingency conditions *I*, at least some laws may apply in the form *if* P *then* Q.

There are recurrent patterns in nature, which often suggest broad generalizations and understanding of phenomena. Strict general laws, like in physics or chemistry, may not be possible in biology, but generalizations, which allow a certain/limited degree of prediction, may be encountered. The problem is that in many cases the lawlike generalization might be decoupled from causality, weakening its power of explanation. Here, I make a case on the importance to study causality to understand the well-known phenomenon of the Species-Area Relationship (SAR), important correlation in the form of a lawlike generalization in ecological theory.

McArthur and Wilson (1967), in the groundbreaking work on Island Biogeography, first formalized the SAR. The correlation is very simple and intuitive: there is a positive correlation between area size and species-richness. Since McArthur and Wilson (1967), the SAR has been showing to be a robust correlation by several experiments, and mathematically deductible to some degree – species-area curves may account for the empirical observation, giving predictive power. The universality of the generalization has been debated, and Storch et al. (2012), for example, argued that the SAR is universal at continental scales. The SAR has been heavily cited and studied since the 1970’s (more than 3.000.000 entries in the scholar google – accessed May.01.2015), and it seems to be one of the triumphs of a robust ecological theory.

Macroevolution and species diversification research have been invigorated during the past 20 years. Advances in molecular biology, phylogenetic comparative studies, and experimental settings have contributed to the progress and broad interest in the field. As a natural consequence, the SAR received an evolutionary interpretation. Losos and Schluter (2000) found positive correlations between area size and *in situ* speciation in *Anollis* lizards on Caribbean islands. Losos and Parent (2010) called the relationship Speciation-Area relationship, giving two empirical examples that fit the model, and emphasizing further generality.

In general, the mechanistic explanations for the pattern remain obscure, and, although authors propose causal explanations to support the different versions of SAR, they are usually not directly tested or studied. The evolutionary SAR may prove to be strong in further testing; nevertheless, because of the nature of evolutionary biology, the pattern may be contingent and constrained to biotic and abiotic factors. Such a claim may suggest the necessity to study causality.

In this article, I suggest a focus on mechanistic explanations on the SAR. If area is an important factor to promote speciation and support species-richness, studying the cause-effect of this phenomena may be explicit, even if there is no single general pattern to explain the cause of the area effect above all group of organisms.

The article is organized as follows: In section 2, I clarify the mechanistic reasoning towards the SAR, focusing on a Darwinian approach to the area effect to exemplify the rationale. In section 3, I develop the idea of the search for associated or secondary patterns in nature in attempt to uncover causality in SAR and interpret the histories of organisms. In section 4, I conclude and summarize my arguments and findings.

## 2 Mechanisms in SAR

### 2.1 Mechanistic approach, evolutionary biology and SAR

In biology, evolution remains in the forefront of understanding of natural phenomena. The famous Dobzhansky’s quote “nothing in biology makes any sense except in the light of evolution” is a strong positioning (Dobzhansky, 1973, p.1). In species diversification, and therefore, in the development of species-richness, there is an ancestral-descendent causal chain between a series of events. Mechanisms, according to Machamer et al. (2000, p.3), “are entities and activities organized such that they are productive of regular changes from start or set-up to finish or termination conditions”. Mechanisms aim to explain how a phenomenon occurred/occurs as a result of the causal chain(s) of its integrated parts. Evolutionary biology and ecology may be explained through a mechanistic approach. The history of the organisms and ecological interactions may be explained through the dissection of a series of events, the interplay of processes, and causal chains. The SAR is a strong pattern recurrent in nature, but as important as to notice the pattern, is to understand exactly what causes it in an evolutionary context.

### 2.2 Darwin’s evolutionary mechanism of the area effect

Darwin in the Origin of Species (1859) offered a possible casual explanation of the area effect in respect to species diversification. The process of change, often unnoticed or underestimated, appears in the chapter on natural selection. For Darwin, selection is a great force in driving evolution, either with or without diversification. He wrote, when discussing the importance of isolation vs. area on generating biodiversity: “although isolation is of great importance in the production of new species, on the whole I am inclined to believe that largeness of area is still more important, especially for the production of species which shall prove capable of enduring for a long period, and of spreading widely”. His reasoning involves population size and complexity of the larger area. Darwin’s mechanistic explanation relies on a series of correlated events as follows: larger areas might have species with larger population size; larger population size might proportionate more variation within the species; with more variation, natural selection might act in different directions - as a result, selection might drive divergence or anagenesis. He also argued that, in larger areas, a species might evolve to a more competitive outcome, spreading to new areas, or simply overcompeting the local species. In addition, larger areas might be more complex in respect to the interactions between species and individuals, altering the selection regime, and potentially generating change and diversity.

These highly intuitive explanations are causal. They address the issue of cause-effect, although they are very much speculative. Nonetheless, some of the claims are pertinent and may deserve contemporary attention in testing their validity.

## 3. Searching for patterns and causality

### 3.1 Integrating, comparing and interpreting patterns

How might one validate causal mechanisms in evolutionary biology and, specifically, in SAR generalization? Patterns are indirect messages that nature sends to us. If there is a pattern of some sort, a causal mechanism might explain what has been demonstrated. In evolution, direct evidence is very scarce, especially those ones which aim to explain historical mechanisms and pathways. In order to have a causal explanation, indirect evidence from different kinds of phenomena may be integrated, compared and interpreted. Secondary or associated patterns may greatly help in explaining a bigger pattern, as in the case of SAR.

As an example of a pattern with a causal explanation, I propose to use the progression rule (Funk and Wagner, 1995). There is a recurrent pattern in the archipelago of Hawaii that, in a given lineage, *in situ* speciation has proliferated basal species on older islands, and derived species on younger islands. Age of taxon and age of island seem to be strongly correlated in some cases (e.g. Cowie and Holland, 2006; Cowie and Holland, 2008). When there is an integration and comparison of the geological history of the archipelago together with the phylogeny of lineages, it becomes clear that the ancestral species migrates to the near youngest island as the chain is being formed. The associated pattern of linear emergence of islands along time allows a pertinent causal explanation. If dispersal ability is not high enough, there is no back dispersal to the older island from the younger, demonstrating a neat correlation island-age vs. lineage-age. In this case, the causal explanation lies on the relationship dispersability and emergence of the new island in elucidating the phylogenetic pattern.

### 3.2 Species formation, barriers to gene flow, and SAR

Barriers to gene flow are ultimately important to species diversification (Sobel et al., 2009). A causal explanation in the evolutionary SAR might account for the development and history of barriers, as well as the complexity of ecological interactions might be studied when the focus is only on species richness in a given location. Area *per se* is never an appropriate causal explanation to proliferation of species: there will be a need for barriers to drive divergence – either/both ecological and/or geographical.

Abiotic and biotic factors might be studied through the search for associated patterns in order to find an appropriate causal explanation. There are many examples of large landmasses with small biodiversity due to a series of factors: deserts are obvious examples. They may have large areas compared to many small spots on the tropical region; still, they have a small biodiversity because of climate and a series of physiological limitations imposed to the lineages. Some species have the ability to adapt, and may even radiate in this inhospitable habitat, emphasizing the evolutionary particularities. The SAR, evolutionary or otherwise, might be a strong pattern, but its causality must be studied explicitly.

## Conclusions

The problem correlation/causation is very old in science. The SAR is one of the most important correlations in biology. It has been studied for decades and proved to be strong and practical. I suggest that explicitly mechanistic explanations should follow SAR documentation and testing. Also, by comparing the studies, recurrent causal chains might be found for specific lineages in large areas. SAR might have more explanatory power and predictability. In order to manipulate the study system, it is easier if the causal mechanism is known. Area *per se* might not account for causality in species diversification or absolute species richness in larger regions. To identify causal chains, I propose to use the integration, comparison and interpretation of other natural patterns (associated or secondary). Barriers to gene flow, as well as the complexity of ecological interactions might be studied in an attempt to find causality in SAR.

## Acknowledgments

I thank Tudor Baetu for extensive discussion on mechanisms in particular and philosophy of biology in general, for valuable comments and encouragement. I thank Daniel Rubinoff for reading through the text. I thank the Brazilian “Coordenaçâo de Aperfeiçoamento de Pessoal de Nível Superior” (CAPES foundation) for the Doctorate Scholarship.

